## Supplementary materials for "Live-cell RNA imaging with the inactivated endonuclease Csy4 enables new insights into plant virus transport through plasmodesmata"

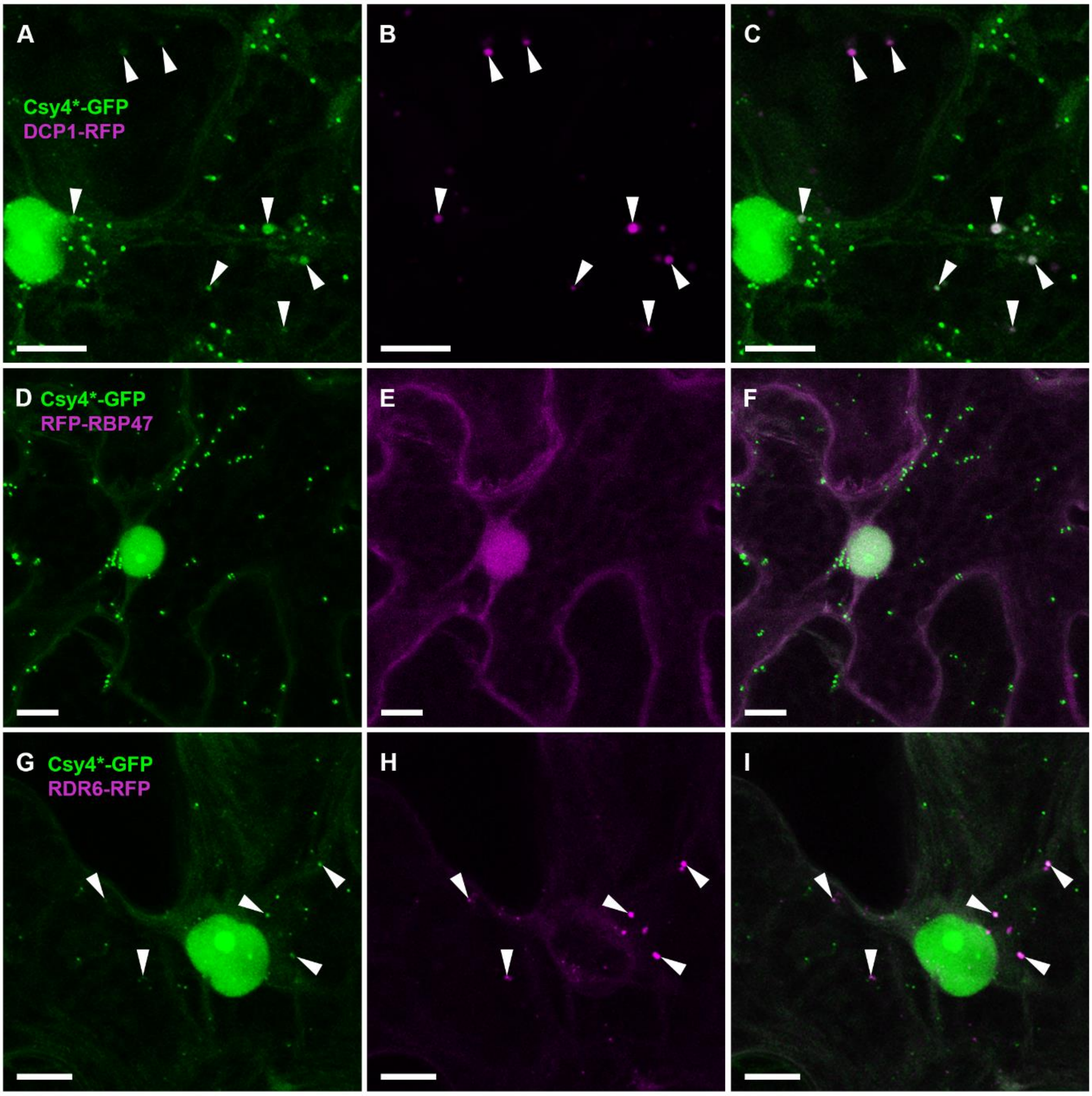


**<u>Supplementary Figure S1:</u> Co-localisation of Csy4*-GFP with different RNA granule markers.** (A) Co-localisation with P-body marker DCP1. Arrow heads indicate DCP1 granules co-localised with Csy4*-GFP. (B) Co-localisation with stress granule marker RBP47. (C) Co-localisation with tasiRNA processing body marker RDR6. Arrow heads indicate DCP1 granules co-localised with Csy4*-GFP. Left column: GFP channel, middle column: RFP channel, right column: merge. GFP fluorescence shown green, RFP fluorescence shown magenta. All images are maximum intensity z-projections. Scale bars, 10 µm.

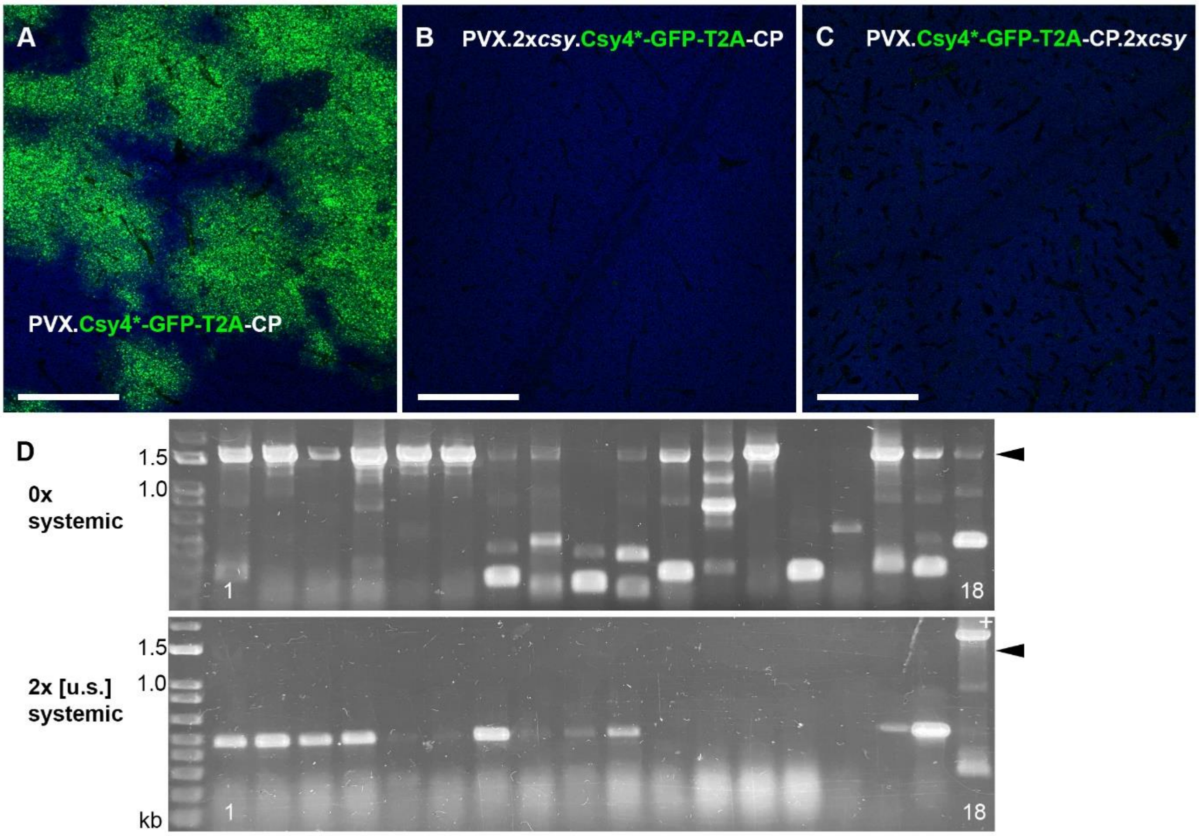


**<u>Supplementary Figure S2:</u> Systemic movement of 'self-tracking' PVX constructs.** (A-C) Representative images from systemic leaves of plants infected with un-tagged PVX.Csy4*-GFP-T2A-CP (A), upstream-tagged PVX.2x*csy*.Csy4*-GFP-T2A-CP (B) and downstream-tagged PVX.Csy4*-GFP-T2A-CP.2x*csy* (C), respectively, at 14 days post inoculation. GFP fluorescence shown green, chlorophyll autofluorescence shown blue. All images are maximum intensity z-projections. Scale bars, 1 mm. (D) RT-PCR analysis of systemically infected leaves at 14 dpi. A segment of the PVX genome from the end of the TGB3 ORF to the beginning of the CP ORF was amplified. Arrow heads on right indicate expected product size when complete Csy4*-GFP-T2A-CP ORF is present. 1 to 18: three biological replicates with six plants each. 0x: un-tagged PVX.Csy4*-GFP-T2A-CP, 2x [u.s.]: upstream-tagged PVX.2x*csy*.Csy4*-GFP-T2A-CP.

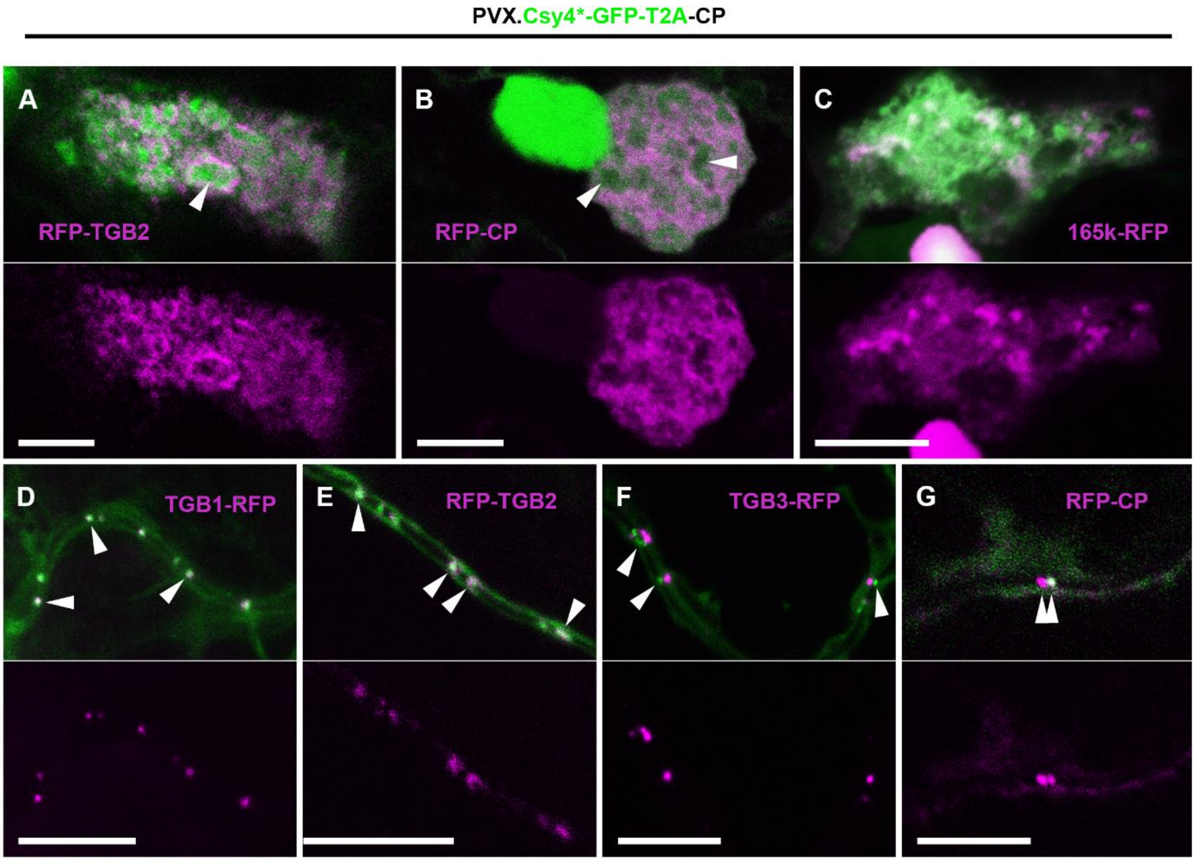


<u>Supplementary Figure S3:</u> **Co-localisations of 'self-tracking' PVX.Csy4*-GFP-T2A-CP with viral proteins expressed *in trans*.** (A) RFP-TGB2 localises to the VRC, appearing to surround vRNA (arrow head). (B) RFP-CP localises to the VRC, also surrounding vRNA (arrow heads). (C) 165k-RFP replicase marker localises to granular structures among the vRNA in the VRC. (D, E and G) TGB1-RFP (D), RFP-TGB2 (E) and RFP-CP co-localise with vRNA inside plasmodesmata (arrow heads). (F) TGB3-RFP is located in membrane structures at plasmodesmata entrances, separate from vRNA inside the channels (arrow heads). Top rows, merged images, bottom rows, RFP channel only. GFP fluorescence shown green, RFP fluorescence shown magenta. All images are single z-sections, except (F) which is a maximum intensity projection of two z-sections. Scale bars, 10 µm.

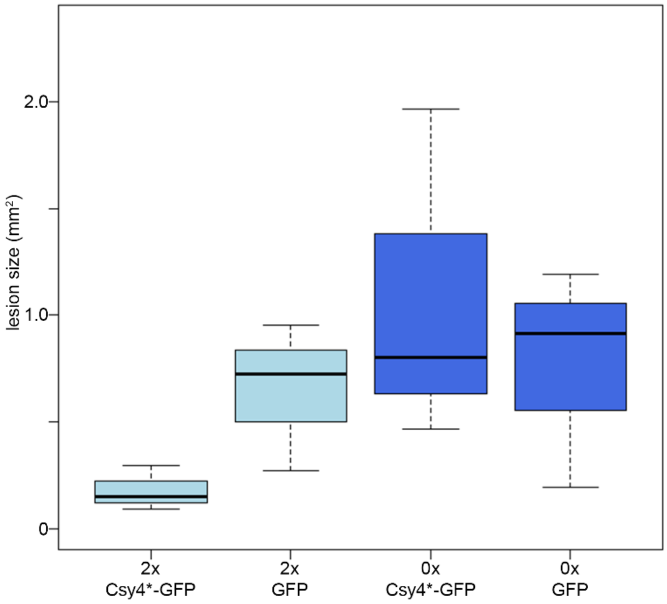


**<u>Supplementary Figure S4:</u> Effect of Csy4*-GFP on PVX spread when expressed *in trans*.** Lesion sizes of untagged (0x) or 2x*csy*-tagged (2x) PVX.RFP-2A-CP at 6 days post inoculation. Virus was inoculated on leaves expressing either unfused GFP or Csy4*-GFP. Two-way ANOVA: The interaction between number of tags and expressed GFP construct was significant (***; $p = 0.000113$). Pairwise Tukey test found no significant differences between treatments.

**Supplementary Table S1:** Oligonucleotide primers used in cloning procedures.

| Primer name | use | Sequence (5'→3') |
|---|---|---|
| DONRfor | sequencing entry clones | CTGGCAGTTCCCTACTCTCG |
| DONRrev | sequencing entry clones | ATGTAACATCAGAGATTTTGAGACACG |
| (Nco)-XT3F | sequencing *csy* tags in PCX | |
| attB-PacDsRF | Gateway cloning of *Pac*I-RFP.2x*ms2*-*Not*I and *Pac*I-RFP.2x*boxB*-*Not*I | AAAAAGCAGGCTGCTTAATTAATGAGGTCTTCCAAGAA |
| M1-DsRR | Gateway cloning of *Pac*I-RFP.2x*ms2*-*Not*I | CCTCATGTATTTACATGGGTAATCCTCATGTTTAAAGGAACAGATG |
| attB-NotM2R2 | Gateway cloning of *Pac*I-RFP.2x*ms2*-*Not*I | AGAAAGCTGGGTGCGGCCGCCACATGGGTAATCCTCATGTATTTAC |
| B1-DsRR | Gateway cloning of *Pac*I-RFP.2x*boxB*-*Not*I | ATTTGCCCTTTTTCAGGGCTTAAAGGAACAGATGGTG |
| attB-NotB2rev | Gateway cloning of *Pac*I-RFP.2x*boxB*-*Not*I and *Pac*I-TMV CP.2x*boxB*-*Not*I | AGAAAGCTGGGTGCGGCCGCCCTTTTTCAGGGCATTTGCCCTTTTTCAGGGCT |
| attB-PacCPfor | Gateway cloning of *Pac*I-TMV CP-*Not*I, *Pac*I-TMV CP.2x*ms2*-*Not*I and *Pac*I-TMV CP.2x*boxB*-*Not*I | AAAAAGCAGGCTGCTTAATTAATGTCTTACAGTATCACTACTCC |
| ΔPacfor | Gateway cloning of *Pac*I-TMV CP-*Not*I | CCCAATAGAGTTAATCAATTTATGTACTAATGC |
| ΔPacrev | Gateway cloning of *Pac*I-TMV CP-*Not*I | GCATTAGTACATAAATTGATTAACTCTATTGGG |
| attB-NotCPrev | Gateway cloning of *Pac*I-TMV CP-*Not*I | AGAAAGCTGGGTGCGGCCGCTCAAGTTGCAGGACCAGAG |
| M1-CPrev | Gateway cloning of *Pac*I-TMV CP.2x*ms2*-*Not*I | CATGTTACATGGGTAATCCTCATGTTCAAGTTGCAGGACCAGAG |
| attB-NotM2rev | Gateway cloning of *Pac*I-TMV CP.2x*ms2*-*Not*I | AGAAAGCTGGGTGCGGCCGCCACATGGGTAATCCTCATGTTACATG |
| B1-CPrev | Gateway cloning of *Pac*I-TMV CP.2x*boxB*-*Not*I | ATTTGCCCTTTTTCAGGGCTCAAGTTGCAGGACCAGAG |
| attB1 adapter | Gateway cloning | GGGGACAAGTTTGTACAAAAAAGCAGGCT |
| attB2 adapter | Gateway cloning | GGGGACCACTTTGTACAAGAAAGCTGGGT |
| sgP25kR | Amplification of PVX vector for TMV 30k insertion | CTTATTCAAATCTCTAAGGTAACTTAACG |
| 2AF | Amplification of PVX vector for TMV 30k insertion | AATTTTGACCTTCTTAAGCTTGCGG |
| sgP25k-30kF | Amplification of TMV 30k for insertion into PVX | aagttaccttagagatttgaataagATGGCTCTAGTTGTTAAAG |
| sgPCP-30kR | Amplification of TMV 30k for insertion into PVX | gtttctgcatctaTTAAAACGAATCCGATTCG |
| 30k-sgPCPF | Amplification of 4x*csy*.RFP-2A-CP with overlap to TMV 30k | ggattcgttttaaTAGATGCAGAAACCATAAG |
| 2A-RFPR | Amplification of 4x*csy*.RFP-2A-CP with overlap to PVX vector | ccgcaagcttaagaaggtcaaaattTCTAGATCCGGTAAGGAAC |
| PVX-Csy4F | Amplification of Csy4-GFP for | tcgaaagaggtcagcaccagctagcATGGACCACTACCTCGAC |

| | | |
|---|---|---|
| | ligation into 'self-tracking' PVX | |
| T2A-GFPR | Amplification of Csy4-GFP for ligation into 'self-tracking' PVX | ccctgccctcgccggagcctccggaCTTGTACAGCTCGTCCATG |
| 3UTR-NotF | Insertion of *Not*I site after CP stop codon | ccgcCTACGTCTACATAACCG |
| 3UTR-NotR | Insertion of *Not*I site after CP stop codon | ccgcTTATGGTGGTGGTAGAG |
| OvercoatF | RT-PCR confirming PVX infections | TTGGCTTGCAAACTAGATGC |
| CP28R | RT-PCR confirming PVX infections | TCCGGGATAGTGAACAGG |
